# Joint cell type identification in spatial transcriptomics and single-cell RNA sequencing data

**DOI:** 10.1101/2023.05.29.542559

**Authors:** Agnieszka Geras, Kacper Domżał, Ewa Szczurek

**Affiliations:** Faculty of Mathematics and Information Science, Warsaw University of Technology, Warsaw, Poland; Faculty of Mathematics, Informatics, and Mechanics, University of Warsaw, Warsaw, Poland; Institute of AI for Health, Helmholtz Center Munich, Neuherberg, Germany

**Keywords:** probabilistic model, MCMC sampling, spatial transcriptomics data, cell types, single-cell RNA-seq data

## Abstract

Understanding the intricate composition of tissues in complex living organisms is crucial for unraveling the mechanisms underlying health and disease. This study addresses the challenge of dissecting cell types within tissues by integrating information from two powerful experimental techniques: single-cell RNA-sequencing (scRNA-seq) and spatial transcriptomics (ST). While scRNA-seq offers insights into transcriptional heterogeneity at the cellular level, ST provides spatial information within tissues. Current methods for cell-type annotation in scRNA-seq and mixture decomposition in ST data are often conducted independently, resulting in reduced statistical power and accuracy. To bridge this gap, we propose ST-Assign, a novel hierarchical Bayesian probabilistic model that jointly performs cell-type annotation in scRNA-seq data and cell-type mixture decomposition in ST data. ST-Assign accounts for shared variables such as gene expression profiles and leverages prior knowledge about marker genes, amplifying statistical strength and mitigating experimental noise. The model’s excellent performance is demonstrated on simulated and real mouse brain data, showcasing accurate cell-type mixture decomposition and cell-type assignment. In comparison to existing tools, ST-Assign demonstrates superior capabilities, particularly in the task of assigning cell types to individual cells. ST-Assign enables exploring the spatial composition of cell types and holds the potential for enhancing our comprehension of diverse biological systems.

## Introduction

Tissues of complex living organisms are composed of millions of cells that come from a large variety of cell types and act in an orchestrated manner (1). The study of individual cell transcriptomes is pivotal to understanding the biological mechanisms behind health and disease (2).

Specifically, single-cell RNA-sequencing (scRNA-seq) is commonly used to examine the extent of transcriptional heterogeneity between cells and identify novel sub-populations within them (3). Cell type annotation is a vital step in the scRNA-seq data analysis pipeline. It is typically conducted through clustering, followed by manual inspection and expert decision-making, which can, unfortunately, be laborious and susceptible to potential biases (4). As a solution, computational tools have been proposed to facilitate automatic, probabilistic cell-type identification, such as CellAssign (5) and scSorter (6).

While scRNA-seq data lack information on cells’ position within the tissue, spatial transcriptomics (ST) technology enables its retention. Nonetheless, the mini-bulk nature of some of popular ST protocols, such as (7) Visium (8) presents the challenge of decomposing hidden cell-type mixtures within each spatial location (spot). To tackle this issue, we have previously developed Celloscope (9), specifically designed to infer the prevalence of each cell type from ST data. Celloscope distinguishes itself from earlier methods for cell type deconvolution in ST spots (10–19) by leveraging prior knowledge of marker genes for cell types. These genes are anticipated to exhibit higher expression in their specific cell types than in other cell types.

When scRNA-seq and ST data originate from the same tissue, an identical set of cell types can be assigned to individual cells in scRNA-seq and deconvoluted in each ST spot. Importantly, variables involved in the data-generating process for these two separate datasets overlap. In particular, marker gene expression profiles characteristic of each cell type rule the observations in both scRNA-seq and ST data. Therefore, a model that would integrate scRNA-seq and ST data based on prior knowledge and such shared variables has the potential to enhance the statistical power of cell-type identification in both data sources. This calls for a novel approach for joint single-cell type identification and cell-type mixture decomposition.

To address this need, here we propose a novel hierarchical Bayesian probabilistic model called ST-Assign, combining signals from two different experimental techniques: (i) single-cell RNA-seq and (ii) spatial transcriptomics. The model offers a computational method to jointly perform the two tasks that until now were carried out independently: (i) cell-type annotation in single-cell RNA-seq data and (ii) cell-type mixture decomposition in spatial transcriptomics spots. In this way, ST-Assign preserves single-cell resolution and, at the same time, benefits from spatial information. Moreover, by extracting the same signal from datasets generated on different platforms, each with distinct yet complementary strengths and weaknesses, the model is less affected by the experimental noise. Importantly, ST-Assign inherits the marker-gene-driven approach to cell type identification from Celloscope (9) and integrates prior domain knowledge about marker genes. This strategy leverages insights derived from multiple independent studies and helps to guide the model inference toward feasible solutions. As a result, ST-Assign effectively analyses unannotated datasets.

We showcase the competitive ST-Assign’s performance on real and simulated data, demonstrating the benefit of probabilistic modeling of combined data sources. ST-Assign enables reliable, in-depth studies of single cells of various types of interest, sequenced both using scRNA-seq and ST.

## Results

ST-Assign takes as input an unannotated scRNA-seq dataset, an ST dataset taken from the same tissue type, prior knowledge about marker genes encoded as a binary matrix, and, optionally, preliminary estimates for the number of all cells within each ST spot (Fig. 1A).

**Fig. 1.**
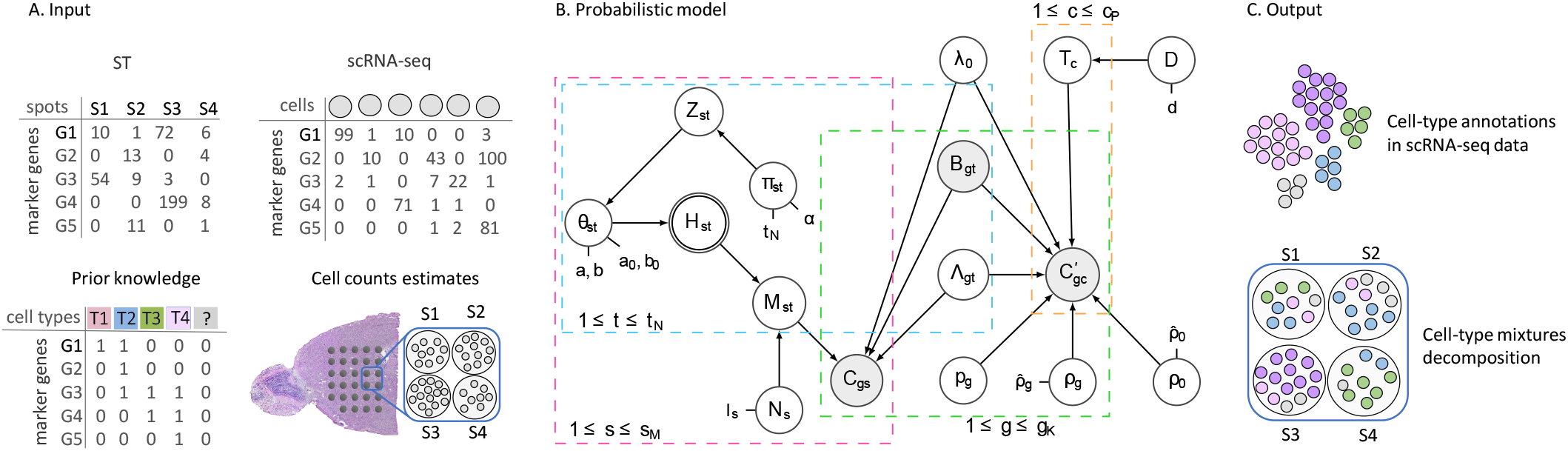
**A**. ST-Assign processes ST and scRNA-seq data, for selected marker genes. The input also includes prior knowledge about marker genes for cell types (including dummy or unknown type, marked as ‘?’) in the form of a binary matrix, along with the total cell count for each ST spot (optional; estimated from H&E image). H&E image source: 10x Genomics. **B**. The graphical representation of ST-Assign. Double-circled nodes correspond to deterministic variables, gray to observed, and the remaining nodes correspond to hidden (inferred) variables. Arrows represent probabilistic dependencies. ST-Assign variables are described in Supplementary Table S1 and its hyperparameters in Supplementary Table S2. Dotted magenta, blue, green, and orange rectangles (plates) represent variables with different indices but identical distributions in the model. **C**. ST-Assign gives as output cell-type mixture decomposition of ST data and annotations for scRNA-seq data.

The probabilistic graphical model behind ST-Assign (Fig. 1B) relies on the expectation that marker genes manifest higher expression in specific cell types across both experiments, facilitating joint decomposition in ST data and cell type assignment in scRNA-seq data. We assume the list of marker genes and their specific cell types are given *a priori*, but gene expression profiles of those cell types are unknown beforehand. As a result, the expression level for each marker gene in each cell type is estimated as a part of the model inference. The model is generative and considers the dependency of the observed gene expression on hidden variables such as the cell type assignment and cell type mixtures’ composition. Moreover, ST-Assign is specifically designed to address the effects of differences between the two experimental techniques by introducing dedicated variables. Conditional probability distributions expressing the dependence structure were used to develop the model’s inference procedure via Markov chain Monte Carlo, specifically the Metropolis-within-Gibbs algorithm.

The model output constitutes cell-type annotations in scRNA-seq data and the count of each cell type in each ST spot (cell-type decomposition; Fig. 1C).

### The study on simulated data proves accurate ST-Assign’s performance

To demonstrate the accurate performance of ST-Assign in both cell-type mixture decomposition and annotation of types to single cells, we executed a study utilizing simulated data. The simulated values of the variables, including variables corresponding to gene expression profiles, crossplatform factors, and the overdispersion parameters, were generated using default priors that ensured similarity to real data (Supplementary Methods). To showcase the excellent performance of ST-Assign regardless of the average number of cells per spot, we investigated scenarios with averages of 5, 10, and 25 cells per spot. Furthermore, to evaluate its robustness to small relative differences between the base expression shared across all genes and the over-expression observed for marker genes, we introduced two options for generating the corresponding values: *basic* (mimicking differences observed in real data) and *lower* (where the average over-expression is five times smaller than in the *basic* scenario; see Supplementary Methods). This nuanced approach allows us to assess the model’s ability to handle realistic variations in gene expression patterns. For each simulation setting, we generated ten replicates of simulated datasets and evaluated the model’s performance in inferring hidden variables. Specifically, to assess the accuracy of estimating the number of cells of each type per spot, we computed the average absolute error across all spots for each replicate *k* using the following equation:

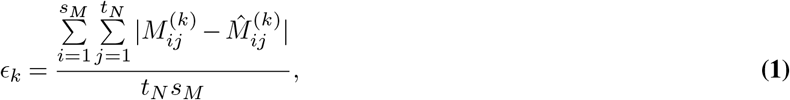

Where 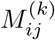 represents the true value of the number of cells of type *j* in spot *i* for the *k*-th replicate, 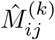 is the number of cells of type *j* in spot *i* for the *k*-th replicate estimated by ST-Assign, *t*_*N*_ is the number of all considered cell types and *s*_*M*_ is the number of spots in ST data. Notably, ST-Assign is the only approach that estimates 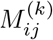 and thus was not compared to any previous method in this task. ST-Assign obtained an excellent average error, with a median below 0.5 cell for 5 and 10 cells per spot and below 2.5 cells for 25 cells per spot (Fig. 2A).

**Fig. 2.**
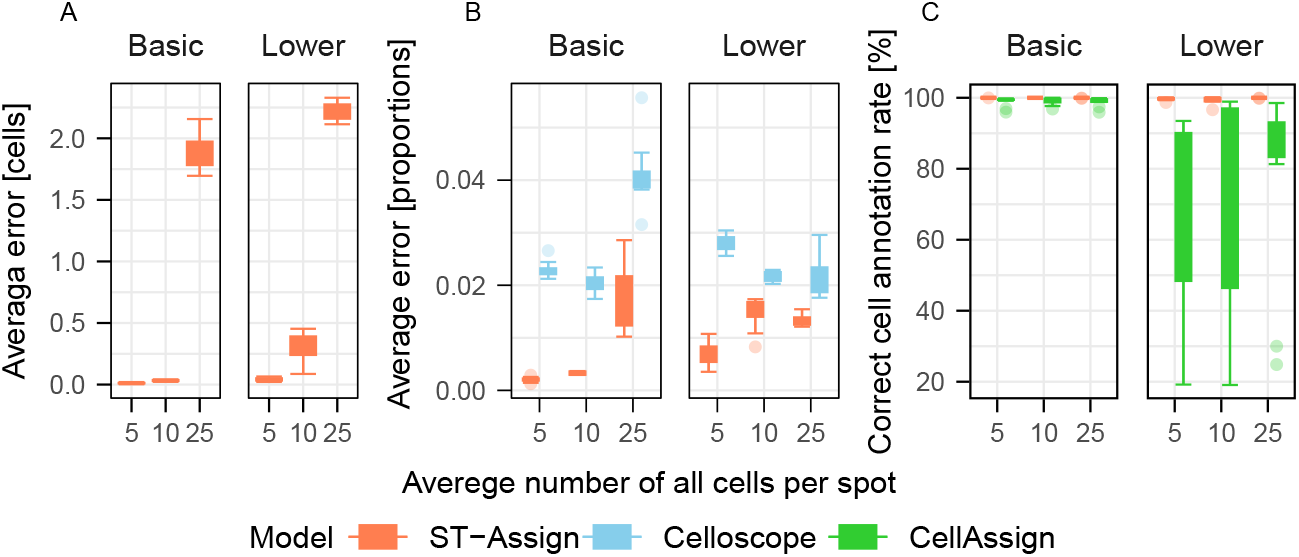
Excellent performance of ST-Assign on simulated data. Box-plots represent distributions of performance values (y-axis) for different methods (colors) and data simulation scenarios (x-axis and panels). **A**. ST-Assign obtains low average absolute error in cell type count estimation; computed using Equation 1, even for large numbers of cells per spot (x-axis). ST-Assign outperformed Celloscope on simulated data in the cell-type mixture decomposition task measured in proportions (**B**) and CellAssign in the cell-type annotation task (**C**).

#### Comparison to Celloscope on simulated data

To assess ST-Assign’s performance against our previous method Cellsoscope (9) (which infers cell-type proportions instead of the number of cells for each cell type), we normalized the *M* vectors to obtain proportions (Fig. 2B). ST-Assign outperformed Celloscope in estimating proportions on simulated data, exhibiting significantly lower errors across all considered data simulation setups. This advantage could be attributed to the stronger statistical signal received by ST-Assign, incorporating information from both ST and scRNA-seq data, a feature not present in Celloscope. Furthermore, it is important to highlight that Celloscope relies on the assumption of the negative binomial distribution for the ST data, utilizing two parameters. In contrast, ST-Assign opts for the Poisson distribution, which requires only one. This choice leads to ST-Assign adopting a simpler approach and a distinction in complexity between the two methods in the context of cell-type mixture decomposition.

#### Cell-type annotation in scRNA-seq data and comparison to CellAssign on simulated data

The excellent performance of ST-Assign was further validated in the task of assigning cell types to individual cells. In this task, ST-Assign demonstrated median accuracy close to 100% for all cells in all the simulation scenarios we considered (Fig. 2C). Importantly, ST-Assign significantly suppressed CellAssign in scenarios characterized by less significant relative differences between base expression and marker gene over-expression (scenario *lower*; Fig. 2C).

### ST-Assign enumerates cells of various types in ST and scRNA-seq mouse brain profiling data

To showcase the performance of ST-Assign in concurrent tasks of ST mixture decomposition and cell type assignment on real data, we leveraged data from two distinct sources: (i) ST data from the anterior mouse brain (sagittal section) from (20) previously analyzed using Celloscope (9), and (ii) a subset of scRNA-seq data from an adolescent mouse brain (21). Specifically, ST-Assign was utilized to determine the number of cells belonging to each type in the ST spots and to assign cell types to individual cells in scRNA-seq data. To determine the set of marker genes for each considered cell type we utilized the set of such genes that were identified as markers by (21) and were also expressed in the ST data (20). The subset of single-cell data (ii) was selected to cover a representative, broad range of cell types and serve as a good illustration of ST-Assign’s capabilities (see Methods for marker gene and cell type selection).

#### Cell type identification

Applying ST-Assign to mouse brain data demonstrated its ability to integrate heterogeneous data sources and facilitate a comprehensive analysis of complex biological systems. The obtained cell type decomposition of ST spots (Fig. 3) is in agreement with previous results acquired with Celloscope on the same dataset, although the previously used gene markers and cell types were a bit different (9). ST-Assign was able to identify known mouse brain structures, effectively differentiate between major neuron types (GABAergic and Glutamatergic), and accurately map the spatial distribution of various cell types, including non-neuronal (such as astrocytes and oligodendrocytes).

**Fig. 3.**
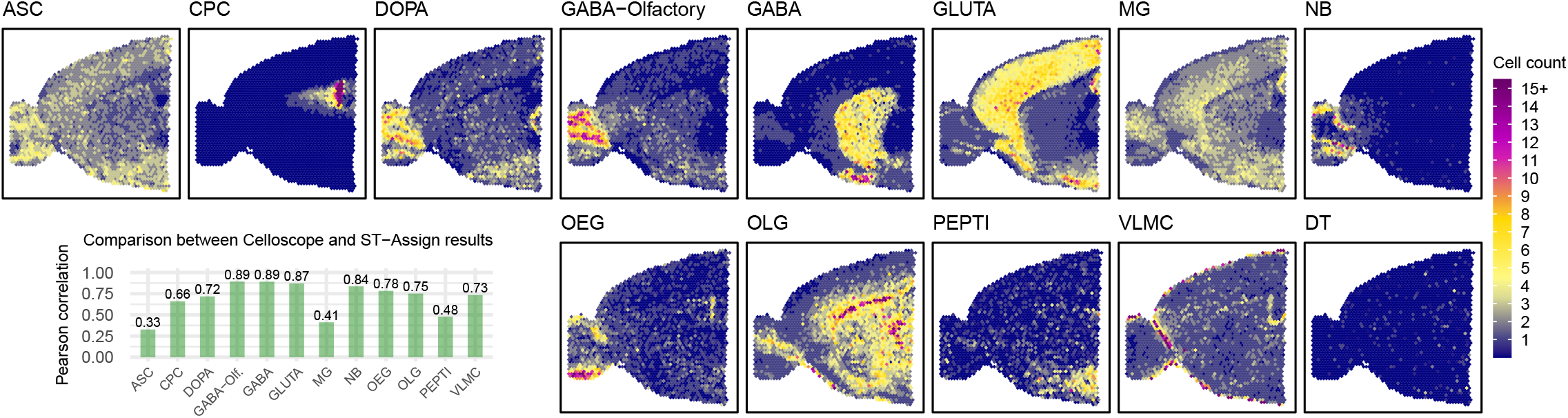
Spatial composition of cell types in the anterior part of the mouse brain (sagittal section) as inferred by ST-Assign. Heatmaps show the abundance of selected cell types, with dark blue indicating absence, yellow moderate occurrence, and magenta dominance. The cell types are as follows: ASC - Astrocytes, CPC - Choroid plexus epithelial cells, DOPA - Dopaminergic neurons, GABA-Olfactory - GABAergic neurons in the olfactory, GABA - GABAergic neurons in the striatum, GLUT - Glutamatergic neurons, MG - Microglia, NB - Neuroblasts, OEG - Olfactory ensheathing glia, OLG - Oligodendrocytes, PEPTI - Peptigeneric neurons, VLMC - Vascular and leptomeningeal cells, DT - dummy type. The barplot displays the Pearson correlation coefficient between results obtained with ST-Assign and Cellocope for each cell type.

As another part of the ST-Assign inference, we performed cell type annotation in the accompanying scRNA-seq dataset (Fig. 4A). To present our findings, we employed an independent t-SNE visualization of the single-cell dataset (Fig. 4). The standard single-cell data analyses pipeline and visualization were carried out using Seurat package (22), incorporating to the dataset marker genes associated with the considered cell types from the binary matrix encoding prior knowledge. The cells were color-coded based on the outcomes of the ST-Assign analysis (Fig. 4A), CellAssign results (Fig. 4B), and annotations sourced from the (21) study, which we regard as the ground truth (Fig. 4C).

**Fig. 4.**
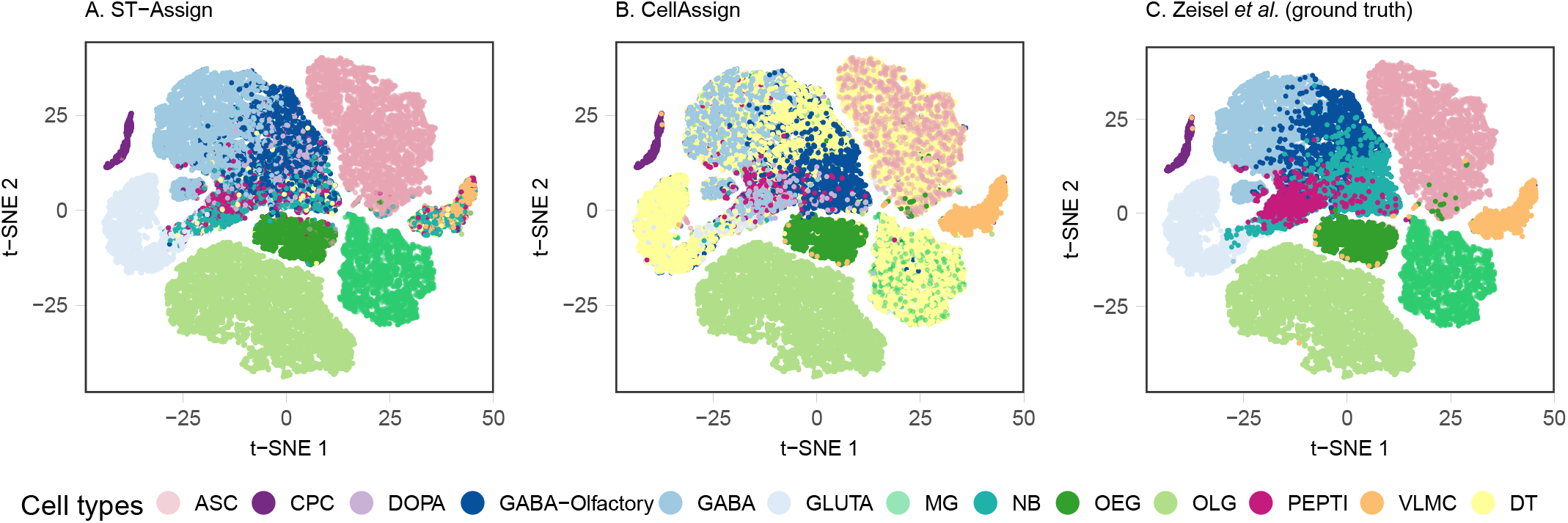
An independent clustering results and t-SNE visualization, with colors indicating the cell type annotation for each cell by **A**. ST-Assign **B**. CellAssign (5), **C**. (21).

The results of ST-Assign (Fig. 4A) generally aligned with the visible scRNA-seq clusters. Moreover, the majority of cells were annotated by ST-Assign to their corresponding cell types (Fig. 5A), as defined by (21). However, a portion of cells (507 out of 38,081) was annotated as the dummy type (Fig. 5A), especially among neuroblasts (204) and vascular and leptomeningeal cells (131), potentially indicating separate sub-types.

**Fig. 5.**
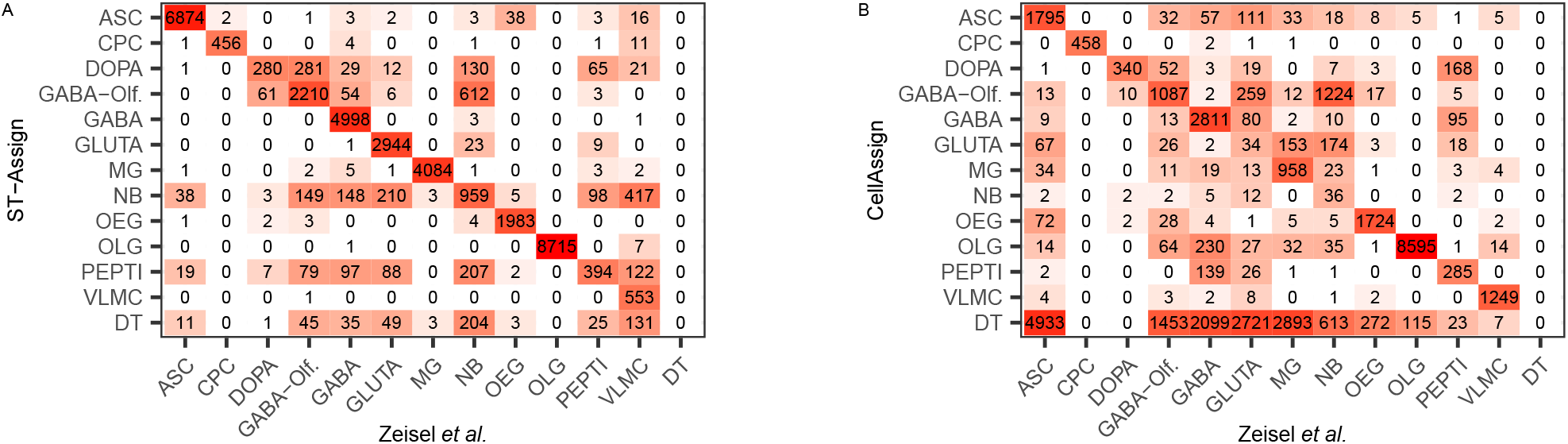
Confusion matrices compare cell type annotations among the ground truth from the study of (21), and ST-Assign (**A**), and CellAssign (**B**). The cell types are denoted by abbreviations, while the counts indicate the number of discrepancies.

#### Comparison to Celloscope on real mouse brain data

To compare the performance of ST-Assign and Celloscope (9), we applied Celloscope on the same mouse brain dataset (Supplementary Fig. S1). The Pearson correlation coefficient was computed to assess the concordance between cell counts estimated by ST-Assign and cell-type proportions estimated by Celloscope (Fig. 3). Overall, a high correlation level was observed, ranging approximately between 0.7 and 0.9. However, specific cell types, such as astrocytes (0.33), microglia (0.41), and peptigeneric neurons (0.48), exhibited lower correlation coefficients. Noteworthy, discrepancies in spatial distribution were evident for astrocytes. For instance, Celloscope showed a low frequency of astrocytes in the olfactory bulb, where ST-Assign indicated their presence. ST-Assign’s results appear more plausible in this context, considering that astrocytes are expected to be distributed throughout the brain.

#### Comparison to CellAssign on real mouse brain data

To compare ST-Assign to its competitor in the task of assigning types to single cells, we applied CellAssign to the same single-cell data (Fig. 4B). CellAssign suffered from both a high rate of assignment of cells to the dummy type (40%) and misclassifications between cell types (Fig. 5B). ST-Assign outperformed CellAssign in correctly identifying glutamatergic neurons and microglia (Fig. 5A). For instance, out of 3,312 cells previously labeled as glutamatergic neurons in (21), 2,721 were unexpectedly annotated as dummy type by CellAssign. For the same cell type, ST-Assign correctly identified 2,944 cells and assigned only 49 of them to the dummy type. In contrast to ST-Assign, the performance of CellAssign was particularly low for astrocytes: 6,946 cells were annotated as astrocytes in (21), CellAssign misclassified 4,933 as dummy type. CellAssign, in turn, exhibited commendable performance for vascular and leptomeningeal cells, aligning with annotations from (21) for 98% of the cells. However, when considering the overall performance, as measured by the Adjusted Rand Index (ARI) with reference to annotations from (21), CellAssign fell short with an ARI of 0.43, while ST-Assign displayed exceptional accuracy with an ARI of 0.93.

## Methods

ST-Assign represents all variables of interest as random variables (Supplementary Table S1). We use a graph-based representation to express the conditional dependence structure between them (Fig. 1B). Since ST-Assign integrates data from the two sources, naturally, the model variables can be divided into three components: (i) variables modeling ST data, (ii) variables modeling scRNA-seq data, and (iii) variables shared between the two components mentioned above.

Let *s* ∈ *𝒮* = {*s*_1_, *s*_2_, …, *s*_*M*_ } index spots, *c* ∈ *𝒞* = {*c*_1_, …, *c*_*P*_ } cells and *g* ∈ *𝒢* = {*g*_1_, *g*_2_, … *g*_*K*_} genes. We assume *t*_*N*_ cell types are present in the considered tissue, indexed by *t* ∈ *𝒯* = {*t*_1_, *t*_2_, …, *t*_*N*_ }. A binary matrix *B* encodes prior knowledge about marker genes, and *B*_*gt*_ takes the value 1 if gene *g* is a marker for cell type *t* and 0 otherwise. We additionally consider a so-called *dummy type*, a helper type that does not have any marker genes. For the dummy type *t*^*dt*^, the value 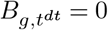 for all marker genes *g*. Consequently, the dummy type gathers all types that have other marker genes than considered *a priori*. Λ_*gt*_ denotes the over-expression of gene *g* when it is a marker for type *t*, and *λ*_0_ represents the base expression profile shared across all the genes. Similarly to our previous model, Celloscope, we assume the increased expression of the marker genes in their specific cell types compared to other cell types. Therefore, we posit that the expected expression measured in a single cell *c* of type *t* equals:

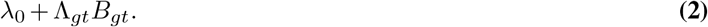

Consequently, if gene *g* is a marker gene for cell type *t*, its expression in cell *c* equals *λ*_0_ +Λ_*gt*_. Otherwise, *B*_*gt*_ = 0, and the expected expression is equal to *λ*_0_. Notably, variable Λ_*gt*_ is shared between the two components modeling ST and scRNA-seq data. Let us notice that ST-Assign considers only a limited subset of genes; therefore, it is easy to check that those considered genes are expressed in the two considered data sets.

ST-Assign serves two purposes: delineating cell-type composition of spots in ST data, represented by the hidden variable *M*_*st*_, and cell-type annotation in scRNA-seq data, defined by the hidden variable *T*_*c*_. Therefore, *M*_*st*_ and *T*_*c*_ are the two central variables in the model. *M* is a matrix with *s*_*M*_ rows and *t*_*N*_ columns. The value of an element *M*_*st*_ is the number of cells in spot *s* that are of type *t*. One row of the matrix *M*, denoted as 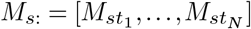 represents the hidden composition of spot *s*. The number of all cells of all types within spot *s* is denoted *N*_*s*_. Clearly, a given row’s entries sum up to *N*_*s*_:

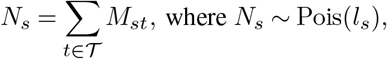

and *l*_*s*_ denotes the preliminary estimates of the number of all cells for each spot (either from counting cells in spots using H&E (23) slides or using a common expected number of cells across spots, given their size).

When it comes to modeling scRNA-seq data, *T*_*c*_ is a categorical variable indicating the cell type of cell *c*:

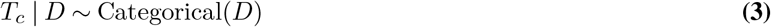

on which we impose Dirichlet prior *D ∼* Dirichlet(*d*) with the concentration parameter *d. D* and *d* are vectors of sizes equal to the number of cell types 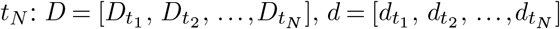.

For gene *g*, its expression in spot *s* denoted *C*_*gs*_, is measured as the count of reads from spot *s* that map to gene *g*. Likewise, the expression of gene *g* in cell *c*, denoted as 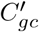, is measured as the count of reads that map to gene *g* from cell *c*. The gene expression matrices *C* and *C*^*′*^ are modeled as observed variables. A row 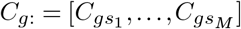 represents the expression of gene *g* across spots and a column 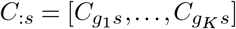 represents the gene expression in spot *s* across marker genes. Anal-ogously, a row 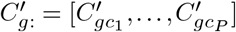 represents the expression of gene *g* across cells and a column 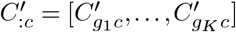 represents expression of cell *c* across marker genes.

Let us denote 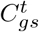 as the number of counts coming from a cell of type *t*. We assume that the random variable *C*_*gs*_ depends on the hidden cell-type composition in spot *s*, modeled as the number of cells of each type in the spot 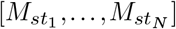. Therefore, *C*_*gs*_ can be expressed as a sum of independent random variables:

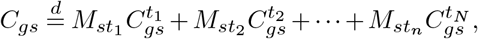

Where 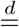 denotes equality in distribution. Therefore:

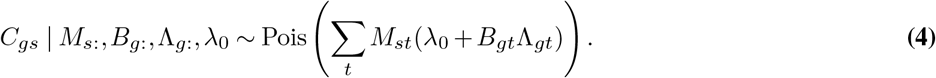

To model *M*_*st*_, we use the multinomial distribution with the number of trials given as the number of all cells in spot s (*N*_*s*_) and the event probability parameter *H*_*s*:_

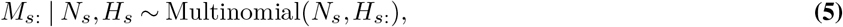

where *H*_*s*:_ is deterministic and computed based on the unnormalized abundance of cell types within spot s:

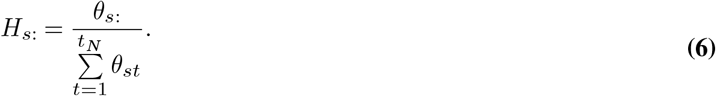

Let us note that *H*_*s*:_ can be interpreted as proportions of cell types within spot *s* and is the primary variable in Celloscope (9). The remaining three parameters, corresponding to the feature allocation model that determines the binary presence of each cell type in each spot: *θ*_*st*_, *Z*_*st*_ and *π*_*st*_ are defined in precisely the same manner as in Celloscope (9), and identical probability distributions are imposed on them (Supplementary Methods).

To model gene expression counts in scRNA-seq data, we employ the negative binomial distribution, where *p*_*g*_ serves as the success probability parameter for gene *g*. This parameter accounts for the over-dispersion in gene expression data and is dependent solely on the gene in question. We posit:

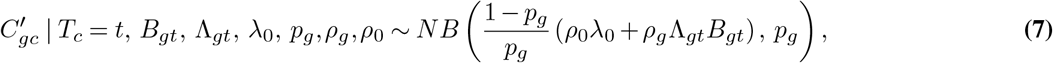

where *ρ*_*g*_ denotes cross-platform factor accounting for differences in sequencing depth between the two techniques. Here, we distinguish between gene-dependent factors, different for each marker gene (*ρ*_*g*_) and a shared factor for non-markers (*ρ*_0_). We assume a Normal prior on *ρ*_*g*_, with the mean 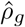estimated from the data:

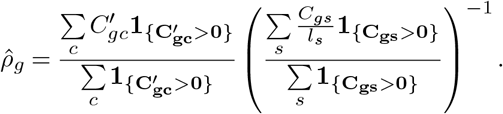

This corresponds to the ratio of the average nonzero expression of gene *g* across single cells in scRNA-seq to the average nonzero expression of gene *g* in ST data. Similarly, a Normal prior is imposed on *ρ*_0_, with the mean computed as

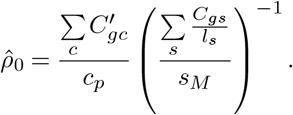

This corresponds to the ratio of the average overall expression of gene *g* across all single cells to the average expression of gene *g* in ST data (including 0 expression). The term 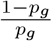 in Equation 7 serves as a rescaling factor, ensuring that the assumption from Equation 2 holds.

To infer the model’s hidden variables, we use the Metropolis-within-Gibbs sampler (Supplementary Methods). Sampling steps (Algorithm 1, Supplementary Methods), the model’s equations for full conditional distributions and proposal distributions used in the implementation are provided in Supplementary Methods.

### Mouse brain data and marker genes selection

We used ST data from a sagittal section of the anterior mouse brain provided by (20) (accessed via `SeuratData` package (24)). We considered 2, 696 spots. Across them, a minimum of 8, on average 108.5, and a maximum of 144 genes had non-zero expression. The minimal total expression per spot was 16, the maximal was 13, 745, and the average was 1, 041.6.

We employed a scRNA-seq dataset that covered all regions of an adolescent mouse brain (21). We downloaded gene expression data from http://mousebrain.org/adolescent/, which also featured information on cell types (clusters) and their specific marker genes. We omitted cells with a total expression below 20, resulting in considering a total of 38, 081 cells. Across cells, a minimum of 4, an average of 28.6, and a maximum of 89 genes had non-zero expression. The maximal total expression per cell was 9, 202, while the average was 307.6.

To define the set of analyzed cell types, we extended the set of cell types selected based on (25), which were previously analyzed in (9) using Celloscope. Here, we included additional types (clusters) from (21). Clusters from (21) that represented brain parts not profiled with ST (e.g., spinal cord) were excluded. The remaining clusters were selected based on the coherent expression of marker genes in ST data (Table 1). A total of 194 genes were chosen and modeled as markers. Marker genes were primarily sourced from (21), supplemented with genes previously by (9).

**Table 1.**
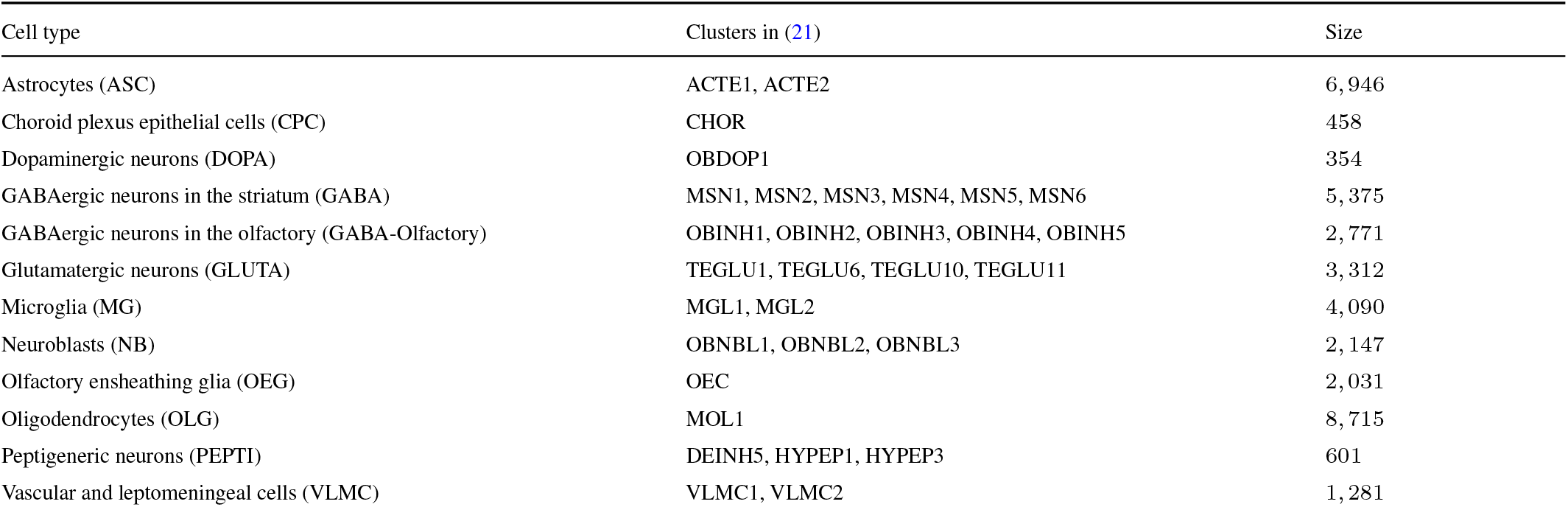
Cell types (their abbreviations in brackets) and their respective clusters in (21) study. In the (21) study, the measured single cells were clustered according to gene expression. These clusters, interpreted as cell types or sub-types, were annotated based on deferentially expressed genes. Columns: analyzed cell types, their respective clusters in (21), and the number of cells in the respective clusters in (21)

The scRNA-seq dataset under examination displayed an elevated percentage of zero entries (approximately 85%) compared to ST data (around 44%). The main reason behind the observed differences in the number of zero entries should be attributed to the fact that the analyzed 10x Genomics Visium ST data consist of *mini-bulk* measurements and as such, may exhibit less technical noise compared to scRNA-seq data. Additionally, this difference may arise from various technical and biological factors, including differences in sequencing depth and biological variability. This justifies using the Poisson distribution to model counts in ST data rather than the negative binomial distribution.

## Discussion

Here, we introduce a novel probabilistic model for concurrent cell-type annotation in scRNA-seq and mixture decomposition in ST data, based on prior knowledge of marker genes. The evaluation carried out on simulated data demonstrated the ST-Assigns’s accurate performance in both tasks. Furthermore, the application of ST-Assign to real mouse brain data showed its utility in the analysis of biological samples.

ST-Assign extends the functionalities of Cellocope (9) to accommodate scRNA-seq data. While leveraging the foundation laid by Cellocope, we have implemented significant modifications, particularly concerning how gene expression given by mixtures of cells in ST data is modeled. Instead of inferring proportions of cell types in each ST spot, modeled as continuous values ranging from 0 to 1, in ST-Assign, we explicitly account for the number of cells (a discrete value) of every type in each ST spot. To achieve this, we utilize the multinomial distribution to model the cell-type composition. Notably, ST-Assign adopts the Poisson distribution to model ST data counts, deviating from Celloscope’s use of the negative binomial distribution. This choice results in a representation with fewer parameters. Importantly, compared to Celloscope, the new model is supplied with additional information on marker gene expression measured in single cells. This new information boosted the performance of inferring gene expression profiles for each cell type. In the task of cell-type annotation, ST-Assign surpassed its rival, CellAssign, as evidenced by its notably higher ARI.

Nevertheless, integrating data from two sources may pose challenges due to technical discrepancies. Cross-platform effects, such as unequal effectiveness with which genes are captured or only partial overlap between sequenced cell types and genes, must be addressed. To this end, we introduced dedicated random variables. Specifically, we added cross-platform factor *ρ* that alleviates unequal sequencing depth, with which genes are measured.

In conclusion, ST-Assign emerges as a valuable tool for the joint analysis of scRNA-seq and ST data, providing a robust solution for simultaneous cell-type annotation and mixture decomposition. Its innovative approach and enhanced capabilities make it a promising asset for understanding complex biological systems at the single-cell level.

## Data and code availability

The ST-Assign’s implementation and data used in this manuscript are freely accessible on GitHub: https://github.com/szczurek-lab/ST-Assign. The results were visualized using ggplot2 (26).

## Ethics approval and consent to participate

Not applicable.

## Supplementary Information

### Supplementary Tables

**Table S1:** Observed, hidden, and deterministic variables in ST-Assign, their corresponding probability distributions, and their interpretations.

| Observed variables |  |  |
| --- | --- | --- |
| $C_{gs}$ | Negative binomial | gene expression for gene $g$ in spot $s$ |
| $C'_{gc}$ | Negative binomial | gene expression for gene $g$ in cell $c$ |
| $B_{gt}$ | - | indicator whether gene $g$ is a marker for cell type $t$ |
| Hidden variables |  |  |
| $\theta_{st}$ | Gamma | unnormalised abundance of cell type $t$ in spot $s$ |
| $\lambda_0$ | - | base gene expression level, shared across all genes and cell types |
| $\Lambda_{gt}$ | - | over-expression for a marker gene $g$ in its specific cell type $t$ |
| $N_s$ | Poisson | number of cells in spot $s$ |
| $\rho_g$ | - | gene-dependent cross-platform factor for gene $g$ |
| $\rho_0$ | - | shared cross-platform factor for non-markers |
| $N_s$ | Poisson | number of cells in spot $s$ |
| $M_{s:}$ | Multinomial | cell type composition in spot $s$ |
| $p_g$ | - | success probability parameter for gene $g$ |
| $Z_{st}$ | Bernoulli | indicator variable stating presence of cell type $t$ in spot $s$ |
| $\pi_{st}$ | Beta | prior for the probability of presence of cell type $t$ in a spot $s$ |
| $T_c$ | Categorical | cell type of cell $c$ |
| $D$ | Dirichlet | prior distribution on $T_c$ |
| Deterministic variables |  |  |
| $h_{st}$ | Dirichlet | fraction of cell type $t$ in a spot $s$ |

**Table S2:**
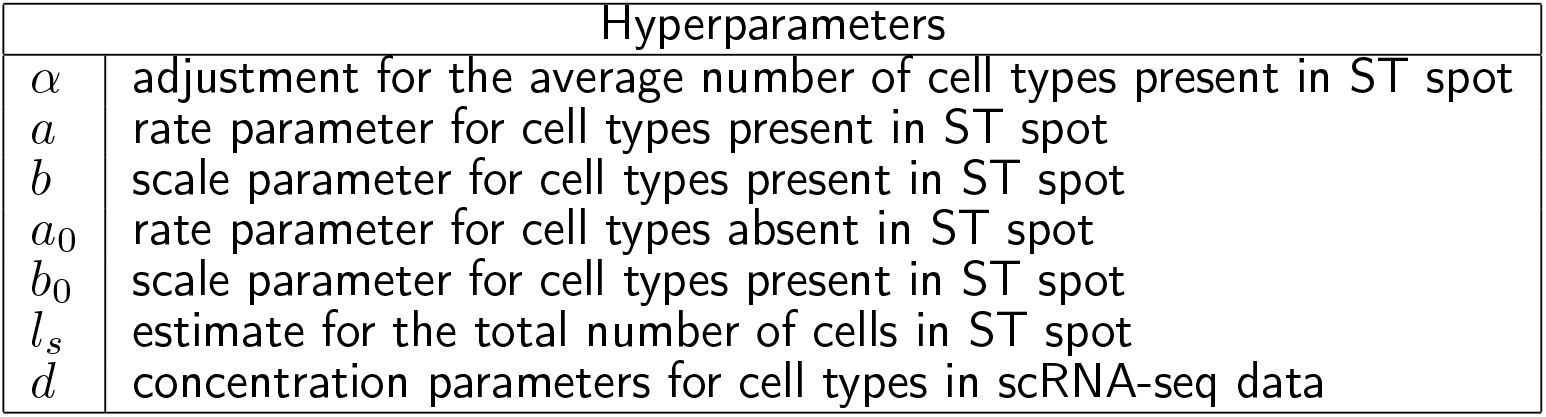
Description of hyperparameters of ST-Assign variables.

### Supplementary Methods

The Metropolis-within-Gibbs sampler is a Markov chain Monte Carlo (MCMC) algorithm which is a combination of the Gibbs sampler and the Metropolis-Hastings method. Suppose the graphical model contains variables *x*_1_, …, *x*_*n*_. Let *MB*(*x*_*i*_) denote the Markov Blanket of *x*_*i*_, i.e., the set containing its parents, children, and co-parents. Then:

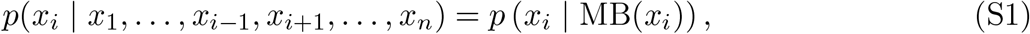

i.e., the conditional distribution of *x*_*i*_ given the values of all other variables equals the conditional distribution given the values of the variables from its Markov blanket.

The iterative sampling procedure is as follows: firstly, starting values for 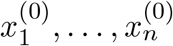 are randomly initialized, then, in a given iteration *j* (*j* = 1, 2, …, *J*, where *J* denotes the number of iterations), we take every variable *x*_*i*_, *i* = 1, …, *n* one by one, in some arbitrary ordering and for each given variable *x*_*i*_, its value is sampled given the values of the variables from the Markov blanket MB 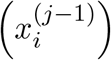 from the previous iteration. As a result each *x*_*i*_ is updated iteratively, up until convergence. There are two options for updating the value of *x*_*i*_ in the *j*-th iteration:

1. If 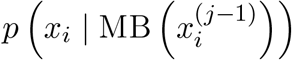 can be expressed in a closed form, a new value 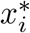 is sampled directly from 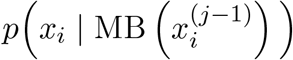and 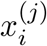is set to 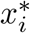.

2. In case we only known a function *g* proportional to *p* (*x*_*i*_ | MB(*x*_*i*_))

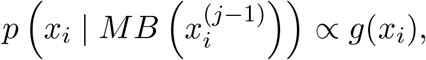

we perform a single Metropolis-Hastings accept-reject step (MH-single-step procedure in Algorithm 1). A candidate value 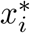 is sampled from a predefined proposal distribution 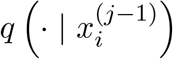, and then either accepted with probability given by

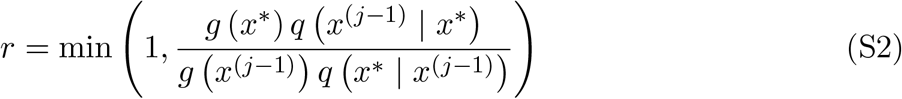

and 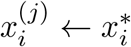, or the previous value is held: 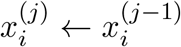. After updating *x*_*i*_, we immedi-ately use the new value for sampling other variables.

#### Algorithm 1 Metropolis-Hastings within Gibbs sampler for inferring ST-Assign parameters

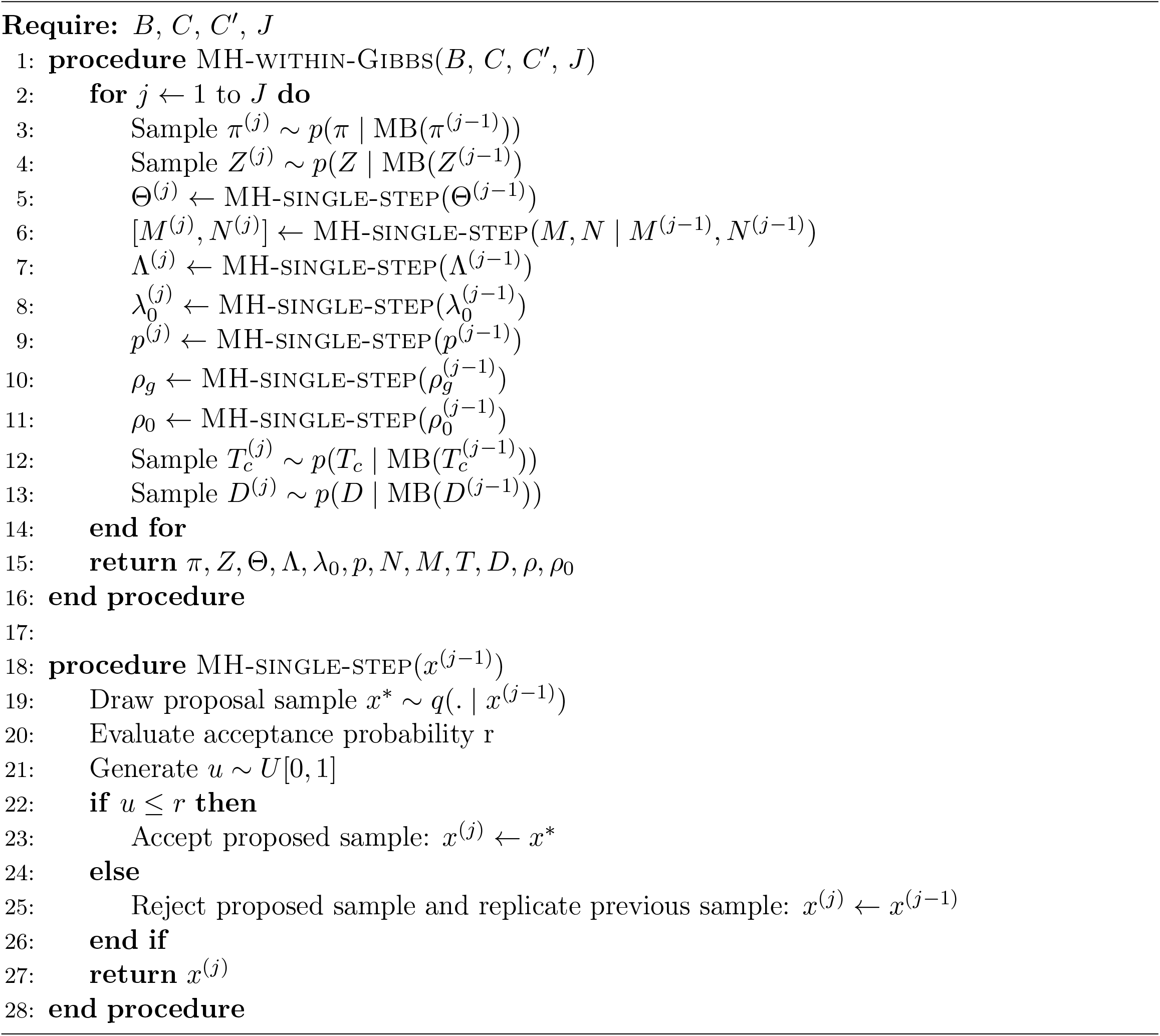

### Inference of the model’s parameters

#### Sampling *π*

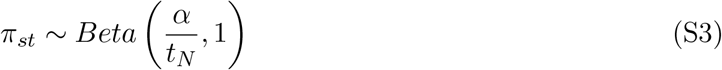

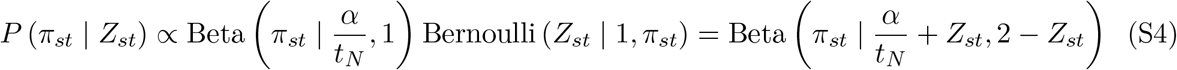

#### Sampling *Z*

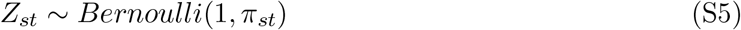

As *Z*_*st*_ is a binary random variable, it suffices to consider its two possible values: 0 and 1.

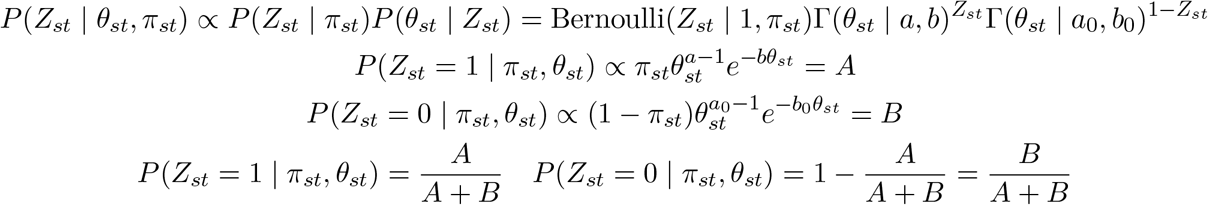

#### Full conditional distribution for *θ*

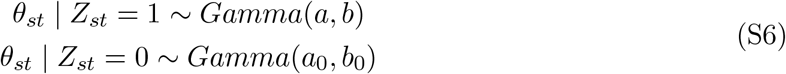

We update each spot independently. For a selected spot *s*:

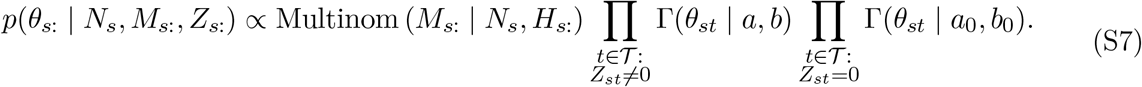

#### Full conditional distribution for *p*_*g*_

For a gene *g*:

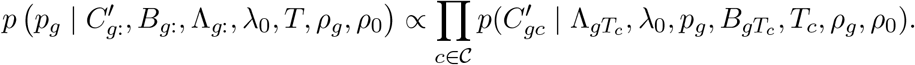

#### Proposal distribution and Hastings ratio for [*N*_*s*_, *M*_*s*_:]

We propose a three-step procedure for proposing new values of *M*_*s*_: and *N*_*s*_. First, to sample a new value 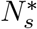 for the variable *N*_*s*_, we choose the ceiling of the truncated normal distribution as the proposal distribution:

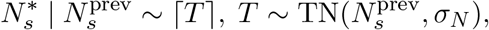

where *σ*_*N*_ controls step size and 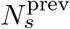 is the value from the previous iteration. Let 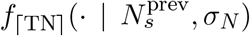 denote the density of ⌈*T* ⌉. Second, we sample a random vector *ϵ ∼ N* (0, *σ*_*M*_ 𝕀), where *σ*_*M*_ controls the step size. Let us denote

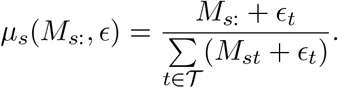

Third, we sample new values 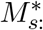of the variable *M*_*s*_: from the multinomial distribution with the number of trials parameter equal to 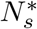 and the event probability parameter computed as a normalized cell-type composition from the previous iteration perturbed with *ϵ*:

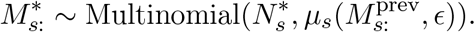

To sum up, the procedure to acquire new values for [*N*_*s*_, *M*_*s*_:] is as follows:

1.Draw a proposal sample 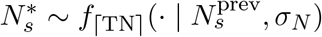.

2.Draw *ϵ ∼* N(0, *σ*_*M*_𝕀).

3.Draw 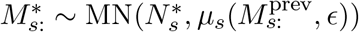.

Finally, the Hastings ratio needs to be computed:

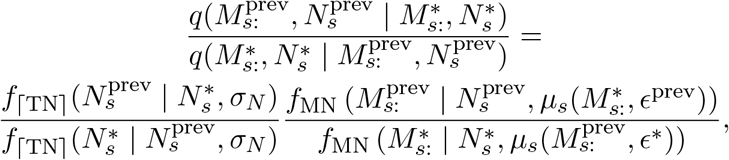

where *f*_MN_ denotes the probability mass function of the multinomial distribution, *ϵ*^∗^ and *ϵ*^prev^ denote the values of *ϵ*, which perturbed the event probability parameter in the current and previous iteration, respectively.

#### Synthetic data simulation

We set *s*_*M*_ = 500 (number of spots), *c*_*P*_ = 2500 (number of cells), *t*_*N*_ = 7 (number of cell types including the dummy type, which corresponds to the last type on the list of types), *a* = 10, *b* = 1, *a*0 = 0.1, *b*0 = 1, *λ*0 = 0.5, *ρ*0 = 1.2, *D* = (1, 1, 1, 1, 1, 1, 0.2) (i.e., the concentration parameter for the dummy type is relatively smaller, as we expect that a minority of cell types is unknown), *α* = 0.5*t*_*N*_ and the total number of marker genes *g*_*K*_ = 160, with their numbers across seven distinct cell types delineated as 11, 15, 20, 27, 37, 50, 0, respectively. To sample gene expression profiles Λ, we first calculate the average gene expression of marker genes for cell types found in a scRNA-seq dataset on the mouse brain cortex [1]. The obtained values were bootstrapped to get values of Λ for all marker genes in the *basic* scenario. To sample Λ in the scenario with *lower* relative difference between *λ*0 and Λ, Λ values were sampled to be, on average, five times smaller than those in the *basic* scenario, while keeping the same *λ*0 values. *ρ*_*g*_ was influenced by the gene expression profiles: genes with higher average expression levels were assigned higher *ρ*_*g*_ values. Additionally, a threshold was applied to ensure the generated *ρ*_*g*_ values remained within a biologically plausible range. The values for *p*_*g*_ were sampled from *Unif* (0, 1), *π* according to Equation S3, *Z* according to Equation S5 and Θ according to Equation S6. We computed *H* based on Θ, from Equation 6 (main text). The values of *M* and *N* were sampled according to Equation 5 (main text), and the mean value of *N* was dependent on the data simulating scenario, each corresponding to the expected total number of cells in each spot, and equal to 5, 10, 25, respectively. *T*_*c*_ was simulated according to the distribution provided by Equation 3 (main text). After all values for parameters have been established, for each of the six considered scenarios, ten replicates were sampled from the generative model: Equation 4 (main text) and Equation 7 (main text). As a result, we obtained 60 synthetic datasets.

## Supplementary Text

In the following section, we outline the run settings that serve for the utilization of ST-Assing, Celloscope, and CellAssign.

### Running ST-Assign

ST-Assign’s run settings were configured as follows: 150,000 iterations were executed, with a burn-in period of 120,000 iterations. The specific parameter values were set as *a* = 10, *b* = 1, *a*_0_ = 0.1, *b*_0_ = 1, and *α* = 5. To determine the initial values and step sizes for the Λ parameter, a data-driven approach was employed. For each gene, the initial value of Λ_*g*_: was set as maximum out of 2 and the average of non-zero entries in *C*_*g*_: divided by 10. The step sizes for Λ parameters were then calculated as one-fifth of these initial values. The step sizes for the rest of the parameters are provided in Table S3. To impose shrinkage prior to the proportion of the *dummy type*, we manually set *Z*_·*t*_ = 0, where *t* here denotes *the dummy type*.

**Table S3:**
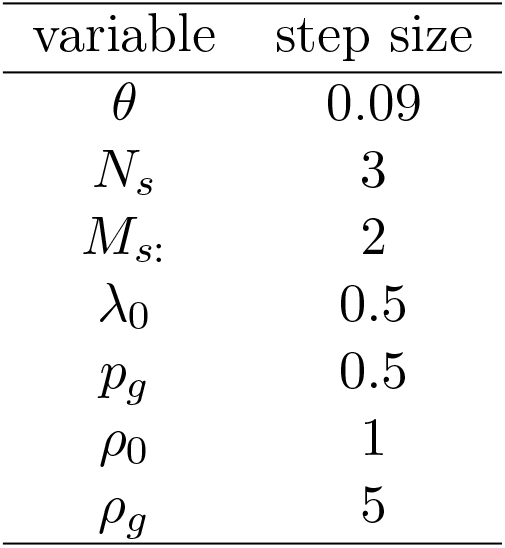
The values of step sizes used for proposal distributions in updating variables through a single accept-reject Metropolis-Hastings step.

### Running Celloscope

The run settings for Celloscope mirrored those used for the sagittal mouse brain in our previous work about Celloscope. However, only one chain was executed (100,000 iterations with 80,000 burn-in period).

### Running CellAssign

We ran CellAssign using the following commands:

~~~
calculated_sum_factors = calculateSumFactors(C_gc)
fit <-cellassign(exprs_obj = C_gc, marker_gene_info = B,
  s = calculated_sum_factors, learning_rate = 10 ^ (−1), shrinkage = TRUE).
~~~

`C_gc` denotes scRNA-seq data. `B` denotes a binary matrix with prior knowledge about marker genes. The learning rate was set to 0.1.

## Supplementary Figures

**Figure S1:**
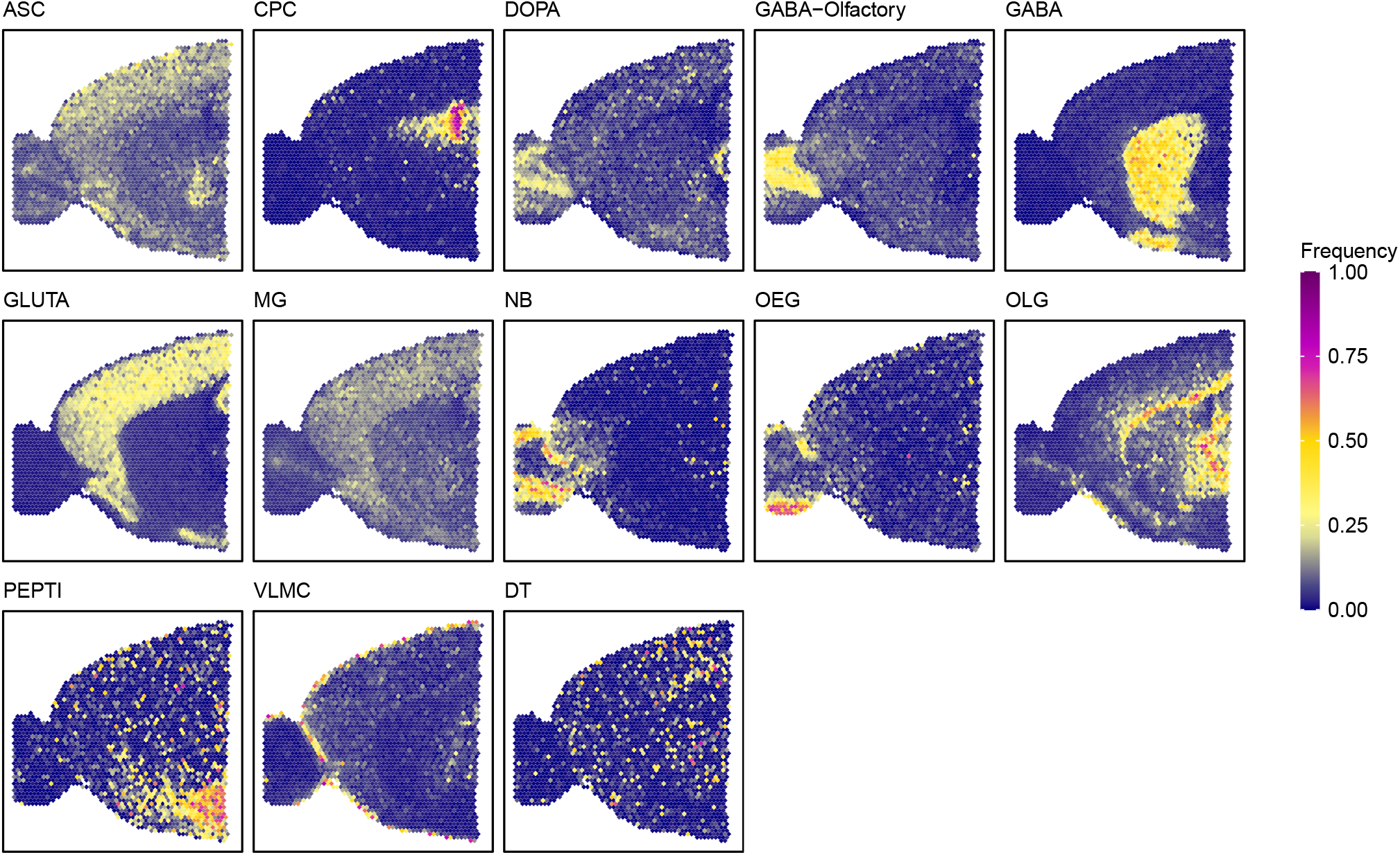
Spatial composition of cell types in the anterior part of the mouse brain (sagittal section) as inferred by Celloscope. Heatmaps show the abundance of selected cell types, with dark blue indicating absence, yellow moderate occurrence, and magenta dominance. The cell types are as follows: ASC - Astrocytes, CPC - Choroid plexus epithelial cells, DOPA - Dopaminergic neurons, GABA-Olfactory - GABAergic neurons in olfactory, GABA - GABAergic neurons, GLUT - Glutamatergic neurons, MG - Microglia, NB - Neuroblasts, OEG - Olfactory ensheathing glia, OLG - Oligodendrocytes, PEPTI - Peptigeneric neurons, VLMC - Vascular and leptomeningeal cells, DT - dummy type.

